# Benchmarking Statistical Multiple Sequence Alignment

**DOI:** 10.1101/304659

**Authors:** Michael Nute, Ehsan Saleh, Tandy Warnow

**Affiliations:** The University of Illinois at Urbana-Champaign

## Abstract

The estimation of multiple sequence alignments of protein sequences is a basic step in many bioinformatics pipelines, including protein structure prediction, protein family identification, and phylogeny estimation. Statistical co-estimation of alignments and trees under stochastic models of sequence evolution has long been considered the most rigorous technique for estimating alignments and trees, but little is known about the accuracy of such methods on biological benchmarks. We report the results of an extensive study evaluating the most popular protein alignment methods as well as the statistical co-estimation method BAli-Phy on 1192 protein data sets from established benchmarks as well as on 120 simulated data sets. Our study (which used more than 230 CPU years for the BAli-Phy analyses alone) shows that BAli-Phy is dramatically more accurate than the other alignment methods on the simulated data sets, but is among the least accurate on the biological benchmarks. There are several potential causes for this discordance, including model misspecification, errors in the reference alignments, and conflicts between structural alignment and evolutionary alignments; future research is needed to understand the most likely explanation for our observations. multiple sequence alignment, BAli-Phy, protein sequences, structural alignment, homology

## Introduction

Multiple sequence alignment is a basic step in many bioinformatics pipelines, including phylogenetic estimation, but also for analyses specifically aimed at understanding proteins. For example, protein alignment is used in protein structure and function prediction [13], protein family and domain identification [49, 23], functional site identification [1, 64], domain identification [5], inference of ancestral proteins [27], detection of positive selection [22], and protein-protein interactions [77]. However, multiple sequence alignment is often difficult to perform with high accuracy, and errors in alignments can have a substantial impact on the downstream analyses [32, 48, 54, 22, 15, 66, 74, 29, 58]. For this reason, the evaluation of multiple sequence alignment methods (and the development of new methods with improved accuracy), especially for protein sequences, has been a topic of substantial interest in the bioinformatics research community (e.g., [74, 69, 28, 56, 34]).

Protein alignment methods have mainly been evaluated using databases, such as BAliBase [3], Homstrad [46], SABmark [73], Sisyphus [2], and Mat-tBench [14], that provide reference alignments for different protein families and superfamilies based on structural features of the protein sequences. Performance studies evaluating protein alignment methods using these benchmarks (e.g., [18, 65, 44]) have revealed conditions under which alignment methods degrade in accuracy (e.g., highly heterogeneous data sets with low average pairwise sequence identity), and have also revealed differences between alignment methods in terms of accuracy, computational efficiency, and scalability to large data sets. In turn, the databases have been used to provide training sets for multiple sequence alignment methods that use machine learning techniques to infer alignments on novel data sets. Method development for protein alignment is thus strongly influenced by these databases, and has produced several protein alignment methods that are considered highly accurate and robust to many different challenging conditions.

An alternative approach to multiple sequence alignment has been developed within the statistical phylogenetics community in which an alignment is co-estimated with a phylogenetic tree by considering stochastic models of evolution in which sequences evolve down a model tree under a process that includes substitutions, insertions, and deletions (jointly referred to as “indels”). Likelihood-based estimation of alignments and/or trees under these models provide a mathematically rigorous and highly appealing approach, and was initially proposed in [6]. Subsequent extensions of this basic approach were made in a sequence of papers [70, 71, 72, 26, 41, 42, 43, 25, 40, 20, 39, 68, 61, 52, 11, 59]. BAli-Phy [68, 61, 59], a Bayesian method that uses MCMC sampling to jointly estimate the multiple sequence alignment and phylogenetic tree under a stochastic sequence evolution model that allows for indels and substitutions, is the most well known of these methods.

Prior studies have shown somewhat different trends with respect to BAli-Phy’s performance on biological and simulated data sets. Three studies [36, 59, 53] evaluated BAli-Phy on simulated nucleotide data sets and found it to have superior accuracy compared to the other alignment methods they examined; this question was examined directly in [36, 59] and indirectly in [53] through the substitution of MAFFT [31] by BAli-Phy within PASTA [44], a divide-and-conquer meta-method that is designed to scale MSA methods to larger data sets. Finally, [30] evaluated BAli-Phy on protein biological benchmarks as well as on simulated protein data sets, and found that BAli-Phy was much less accurate than some other MSA methods (Prank [37], Muscle [19], and variants of MAFFT) on the biological data, but was very good (and for some criteria it was the best) on the simulated data. This study is intriguing but limited, in that they used somewhat non-standard evaluation criteria and did not explore several leading protein alignment methods. In addition, the data sets that were analyzed in [30] were large for BAli-Phy (the simulated data sets had 100 sequences, and the biological data sets ranged up to 100 sequences) and BAli-Phy was only run twice, each for only 1000 MCMC iterations. As discussed in [30, 60], 1000 MCMC iterations may not have been sufficient to allow BAli-Phy to reach convergence on data sets of this size, and it is known that BAli-Phy can have reduced accuracy if stopped prematurely [60]. Hence, a more careful evaluation of BAli-Phy is necessary to understand its performance on biological benchmark data sets.

In this paper, we report on an extensive performance study in which we compare BAli-Phy version 2.3.8 to a collection of leading protein sequence alignment methods. We use 1192 data sets from four established benchmark databases of protein multiple sequence alignments (BAliBASE, Sisyphus, MattBench, and Homstrad) as well as 120 simulated data sets in order to characterize the relative and absolute accuracy of the alignment methods we explore. We limit our study to biological sequence data sets with at most 25 sequences and to simulated data sets (under 6 model conditions) with 27 sequences, so that we are able to run BAli-Phy for long enough to enable it to converge. In particular, we ran BAli-Phy on each data set using 32 independent runs, each for 48 hours (i.e., BAli-Phy was run somewhat longer than 2 months on each data set). This analysis protocol enabled BAli-Phy to generate many hundreds of thousands (and in several cases more than 1,000,000) of MCMC samples for each data set that it analyzed, and achieve good ESS values that indicate that BAli-Phy may have converged well on these data sets. Our study used more than 230 CPU years for the BAli-Phy analyses alone, and provides a careful evaluation of how BAli-Phy performs on biological and simulated data sets.

The most important outcome of our study is that BAli-Phy is dramatically more accurate than all the alignment methods we explore on the simulated data sets, but is among the less accurate on the biological data sets. One possible explanation is that many of the reference sequence alignments in these benchmark data sets have substantial error. Another potential explanation is model misspecification, so that the sequence evolution models underlying BAli-Phy may be a poor fit to how protein sequences actually evolve. Finally, it is possible that many of the reference alignments in the biological benchmark data sets reflect shared structural features that are not a result of descent from a common ancestor (i.e., the aligned amino acids in the reference alignments are structurally homologous and not evolutionarily homologous). Further research is needed to determine the major causes for the discordance between performance on biological benchmarks and simulated data sets.

## Materials & Methods

### Alignment Methods

We explored the following multiple sequence alignment methods: BAliPhy v. 2.3.6, Clustal-Omega v. 1.2.4 [65], CONTRAlign v. 1.04 [16], DiAlign v. 2.2.2 [47, 24], KAlign v. 2.04 [33], MAFFT v. 7.305b [31], Muscle v. 3.8.31 [19], Prank v. 140603 [38, 37], Prime v. 1.1 [78], ProbAlign v. 1.4 [63], Probcons v. 1.12 [17], PROMALS3D [57], and T-Coffee v. 11.00.8cbe486 [50, 51, 55], We explore two ways of running MAFFT: MAFFT-G-INS-i and MAFFT-Homologs (using the SwissProt Database [4]).

All methods other than BAli-Phy and Promals3D were performed in default mode. Promals-3D enables structural alignment features, but we turned these off using the following sample command:

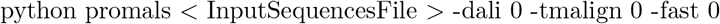

BAli-Phy requires specific parameters (including the substitution model and the number of MCMC iterations) to be set by the user. To select a protein sequence evolution model for use in BAli-Phy, we applied RAxML [67] version 8.2.9 to the alignment computed using MAFFT L-ins-i. We ran 32 independent runs of BAli-Phy, each for 48 hours, discarding the first 25% of the alignments that were generated during the MCMC run, and then retaining every 10th alignment in the remaining sample. The point estimates of the alignments were computed using the posterior decoding (PD). According to the output from BAli-Phy, the vast majority of the BAli-Phy runs we performed showed evidence of having converged, as indicated by various statistics (e.g., Minimum ESS values); see Supplementary materials for these statistics.

### Computational Resources

BAli-Phy and T-Coffee are the most computationally intensive methods we explored, and so these were run on the Blue Waters supercomputer at the National Center for Supercomputing Applications (NCSA); all other methods were run on the Campus Cluster at the University of Illinois at Urbana-Champaign.

### Evaluation Criteria

The accuracy of the estimated alignment was assessed in comparison to the reference alignment for the biological data sets, and to the true alignment for the simulated data sets. We used FastSP v. 1.6.0 [45] to calculate alignment accuracy with respect to the Modeler Score and SP-score. These accuracy measures produce a value between 0.0 and 1.0, with 1.0 indicating perfect accuracy and 0.0 indicating complete failure. We also report the expansion ratio, which is the ratio of the numbers of sites in the estimated alignment and the reference or true alignment; values below 1.0 represent over-alignment (i.e., shorter alignments than the reference or true alignment) and values greater than 1.0 represent under-alignment.

### Data Sets

#### Protein biological data sets

We took all the alignments from the four databases we selected (BAliBASE, MattBench, Homstrad, and Sisyphys) that had between 4 and 25 sequences. All alignments with more than 25 sequences were then sub-sampled to produce a data set with between 5 and 25 sequences; see Supplementary Materials for the protocol used for subsampling.

T-COFFEE failed to align a number of data sets due apparently to a lack of results from the BLAST step of the algorithm; this was particularly pronounced on the BAliBase data, where 82 out of 742 alignments were not completed, although it also failed to align 2 datasets each from the other three benchmarks.

BAli-Phy was able to analyze all the datasets, but on two datasets the posterior decoding algorithm failed due to the high computational complexity of having a small number of very long sequences. After eliminating the datasets where T-Coffee and the Bali-Phy posterior decoding failed to complete, we still had a large number (1192) of reference alignments from the four benchmarks.

Table 1 presents empirical properties for the reference alignments for these 1192 data sets, including average pairwise sequence identity (PID), average sequence length, average number of sequences, average percentage gapped, and mean gap length.

**Table 1:**
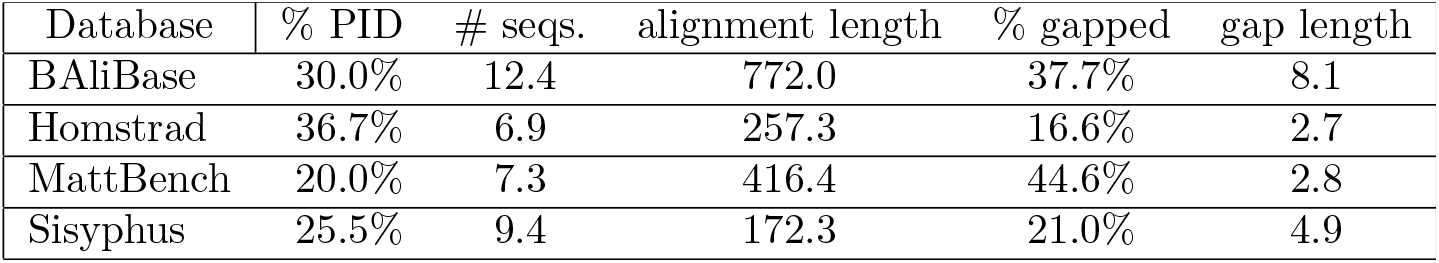
Empirical properties of the 1192 reference alignments from the four biological benchmark collections. We report the average pairwise sequence identity (%PID), average number of sequences, average alignment length, average fraction of the reference alignment occupied by gaps, and median gap length.

| Database | % PID | # seqs. | alignment length | % gapped | gap length |
| --- | --- | --- | --- | --- | --- |
| BAlibase | 30.0% | 12.4 | 772.0 | 37.7% | 8.1 |
| Homstrad | 36.7% | 6.9 | 257.3 | 16.6% | 2.7 |
| MattBench | 20.0% | 7.3 | 416.4 | 44.6% | 2.8 |
| Sisyphus | 25.5% | 9.4 | 172.3 | 21.0% | 4.9 |

#### Simulated data sets

We generated 120 simulated data sets (20 data sets from each of 6 different model trees) to evaluate the alignment methods for this study. To obtain the basic model tree topology and branch lengths, we selected the 27-sequence serine protease data set from the Homstrad benchmark collection, computed a MAFFT L-ins-i alignment on the data set, and then used RAxML to construct a phylogenetic tree with branch lengths. We set the indel rate and the gap length distribution (a negative binomial) to match the empirical distribution for the serine protease data set. We then modified this basic model tree in two ways – by rescaling the branch lengths (by a factor of three) and reducing the indel rate – to produce six different model conditions (Table 2) that ranged in terms of the average percent gapped (from 18.3% to 46.4%) and average pairwise sequence identity (PID) (from 10.7% to 23.9%). Hence, this process produced six different model conditions with a range of average PID and percentage gapped that cover the characteristics of the biological benchmark data sets we explored. The root sequence had 200 amino acids, and sequences evolved down each model tree with substitutions and indels under the WAG [75] model, using Indelible [21].

**Table 2:**
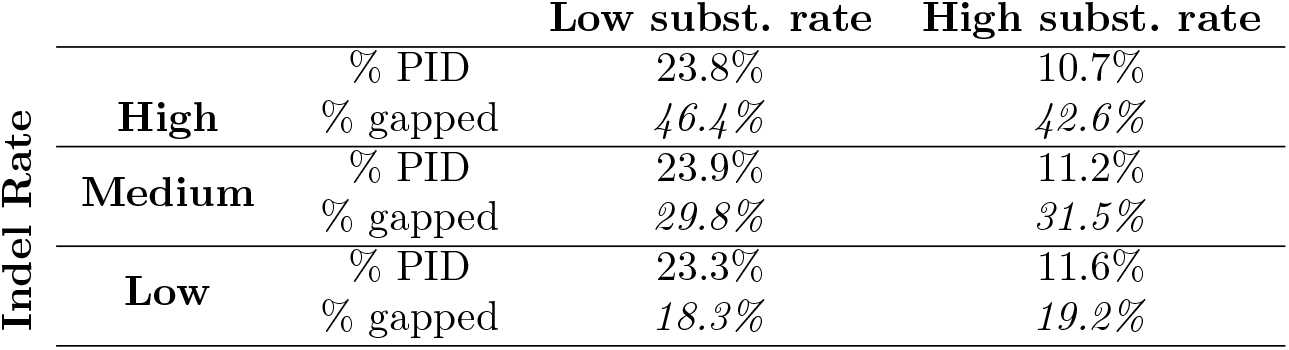
Empirical properties of the true alignments for the simulated data sets, each with 27 sequences. Each submatrix represents one of the six model conditions, and the top row within each submatrix represents the mean percent pairwise identity (% PID) and the bottom row represents the percentage gapped.

## Results

### Results on Biological Data Sets

The results shown are restricted to the 1192 datasets where all methods ran successfully.

Our first experiment examined the overall accuracy of the different methods we examined, showing SP-Score, Modeler Score, and expansion ratio (Fig. 1). Overall, BAli-Phy had the best average Modeler score but among the lowest average SP-score of all the methods. T-Coffee and Promals had the best overall SP-scores, and (except for BAli-Phy) the best Modeler scores. CONTRAlign and MAFFT-homologs were next best, followed by ProbAlign and Probcons. MAFFT-G-ins-i, Prime, Clustal-Omega, and Muscle, roughly in that order, came in the next group. Finally, Prank, Di-Align, and KAlign were the least accurate on these data.

**Figure 1:**
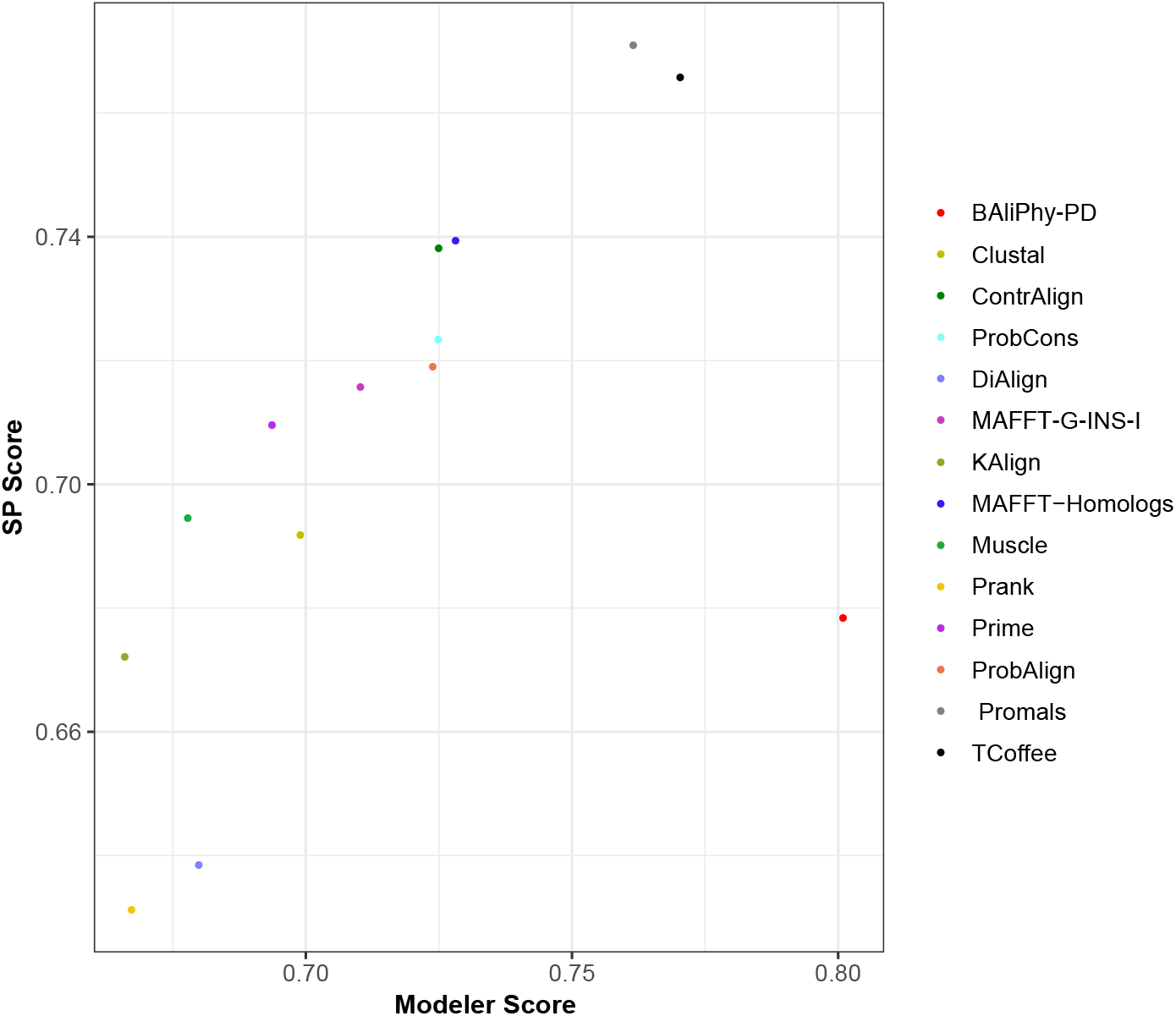
Average Modeler Score vs. average SP-score of the full set of multiple sequence alignment methods on the biological benchmark data sets, each with at least 4 and at most 25 sequences; each data point represents analyses of 1192 data sets from the four benchmark collections (658 from BAliBase, 231 from Homstrad, 202 from MattBench, and 101 from Sisyphus).

The same comparison was performed on the different benchmarks individually, restricted to the set of top-performing methods (i.e., with Prank, Di-Align, and KAlign removed), and those data sets on which all methods completed. While the relative and absolute performance varied to some extent between benchmark collections, BAli-Phy consistently had very good Modeler scores and very poor SP-scores (Fig. 2). T-Coffee and PROMALS were also typically the top two most accurate alignment methods, and Muscle and Clustal-Omega were among the least accurate alignment methods. Alignment accuracy was highest on the Homstrad database (with the average SP-score and Modeler scores nearly always above 80% for all methods), relatively high for BAliBASE, lower for Sisyphus, and lowest for MattBench. Thus the different benchmarks present different levels of difficulty, with MattBench the hardest and Homstrad the easiest.

**Figure 2:**
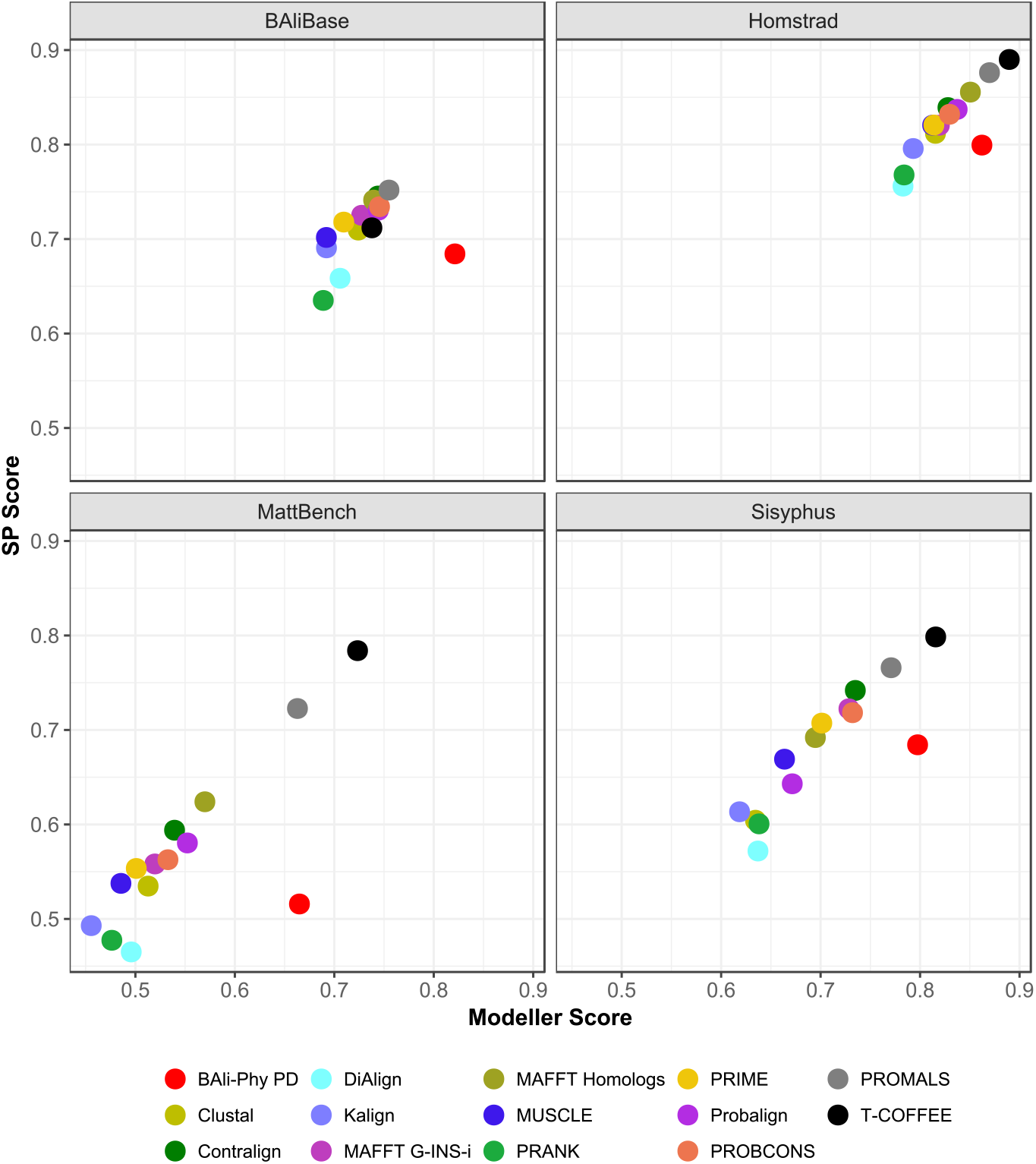
Average Modeler Score vs. SP-Score of the top methods on the biological benchmark data sets, each with at most 25 sequences. Results shown are for 1192 data sets from the four benchmark collections (658 from BAliBase, 231 from Homstrad, 202 from MattBench, and 101 from Sisyphus)

We then examined the impact of average pairwise sequence identity (PID) on alignment accuracy, measured using Modeler score, SP-score, and expansion ratio. Figure 3 shows that when the data sets have sufficiently low heterogeneity (i.e., average PID above 25%), most methods have expansion ratios that are close to 1.0, and so produce alignments that are approximately the correct length. However, when the data sets have high heterogeneity, the only methods that consistently come close to producing alignments of approximately the same length as the reference alignment are Probalign, Probcons, Promals, and T-Coffee. Of the remaining methods, BAli-Phy, Di-Align, and Prank under-align, and the others over-align. Also, BAli-Phy displays the largest degree of under-alignment, and even under-aligns on the bin with the lowest average PID.

**Figure 3:**
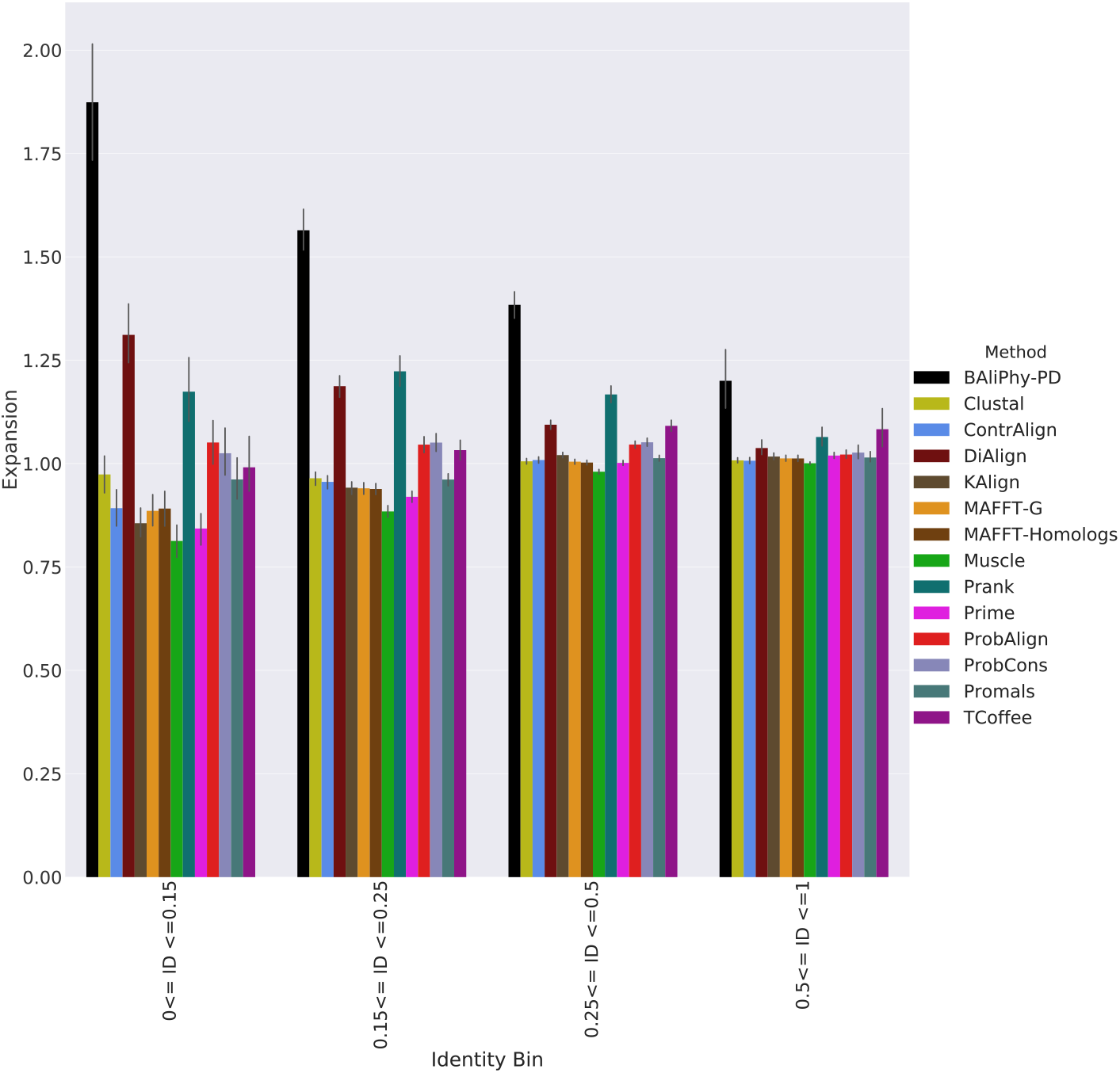
Average expansion ratios on the 1192 biological benchmark data sets, each with at most 25 sequences, by average percent ID (ID). Values more than 1.0 indicate under-alignment (i.e., longer alignments than the reference alignment), while values less than 1.0 indicate over-alignment (i.e., shorter alignments than the reference alignment). The four bins based on average sequence identity, ordered from smallest to largest, have 83, 417, 615, and 77 alignments, respectively.

As seen in Figure 4, the average PID is correlated with the Modeler Score of estimated alignments for all methods, with the best accuracy obtained for the data sets with the highest average percent identity (PID). In addition, the difference between methods is less for the high PID data sets and then increases as PID drops. In particular, for the lowest average sequence identity data sets, there is a big gap between the least accurate methods (Muscle and Clustal-Omega) and the most accurate methods (BAli-Phy and T-Coffee). In addition, while the Modeler score for BAli-Phy is impacted by PID, the impact seems less than for other methods, as BAli-Phy’s Modeler score generally remains high as the average PID is reduced. Also, while T-Coffee clearly ties for best on the lowest average PID data sets, it is not among the best for the highest average PID data sets (and indeed it is the least accurate of the collection). Similarly, Promals clearly dominates all methods other than BAli-Phy and T-Coffee for the lower average PID data sets, but is not noteworthy on the highest PID data sets.

**Figure 4:**
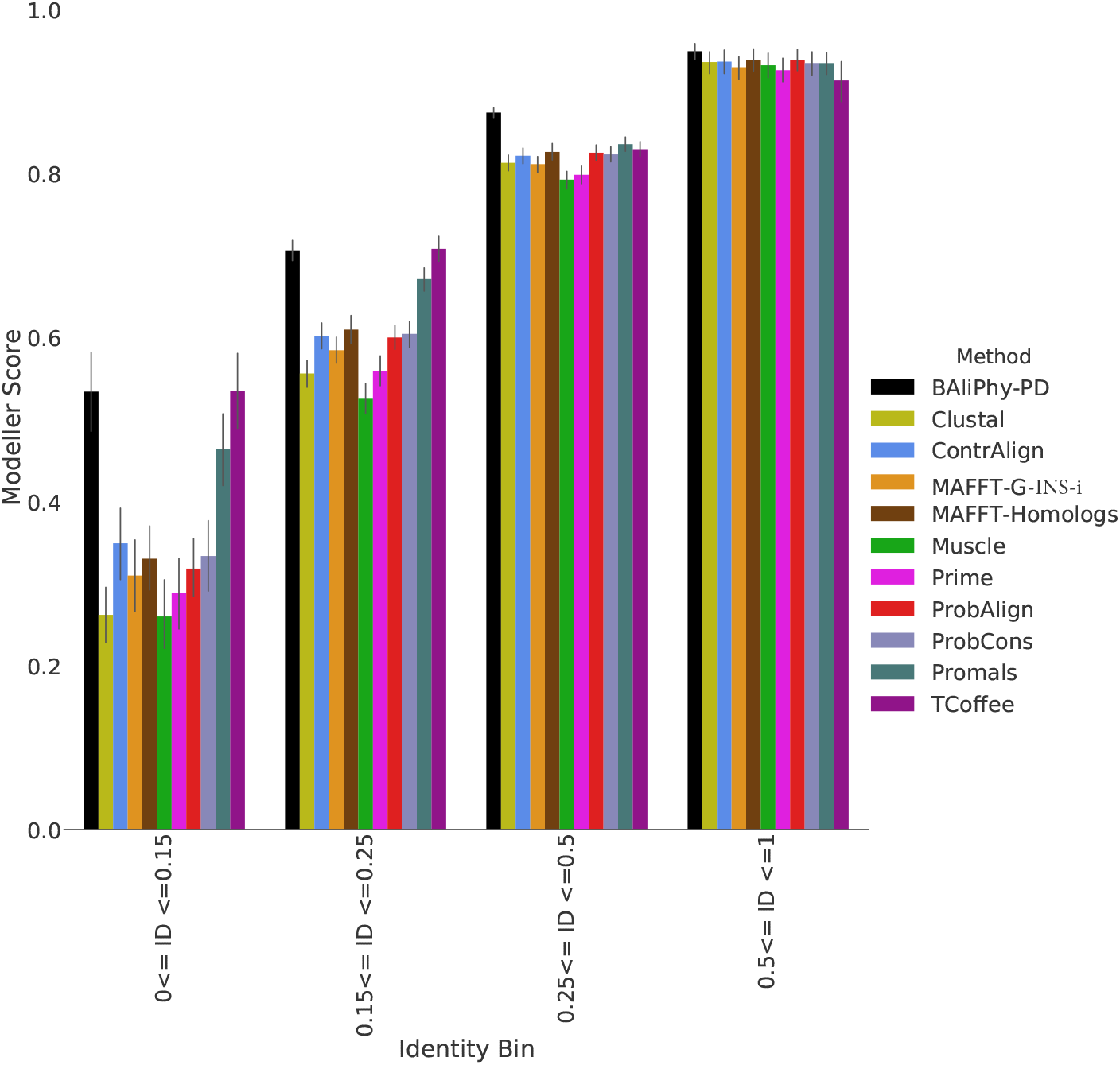
Average Modeler Scores for the top methods on the 1192 biological benchmark data sets, binned into different average pairwise sequence identity (ID) levels. The four bins based on average sequence identity, ordered from smallest to largest, have 83, 417, 615, and 77 alignments, respectively.

Figure 5 enables the same comparison but with respect to SP-score. With the exception of BAli-Phy’s performance, all the trends observed for the Modeler score hold for the SP-score. The best SP-scores are obtained by T-Coffee and Promals, two methods that rely on external biological information, but even the vanilla methods (e.g., MAFFT G-ins-i) are substantially more accurate than Prank and BAli-Phy. Indeed, BAli-Phy is among the worst for SP-score of these top methods, under all tested conditions.

**Figure 5:**
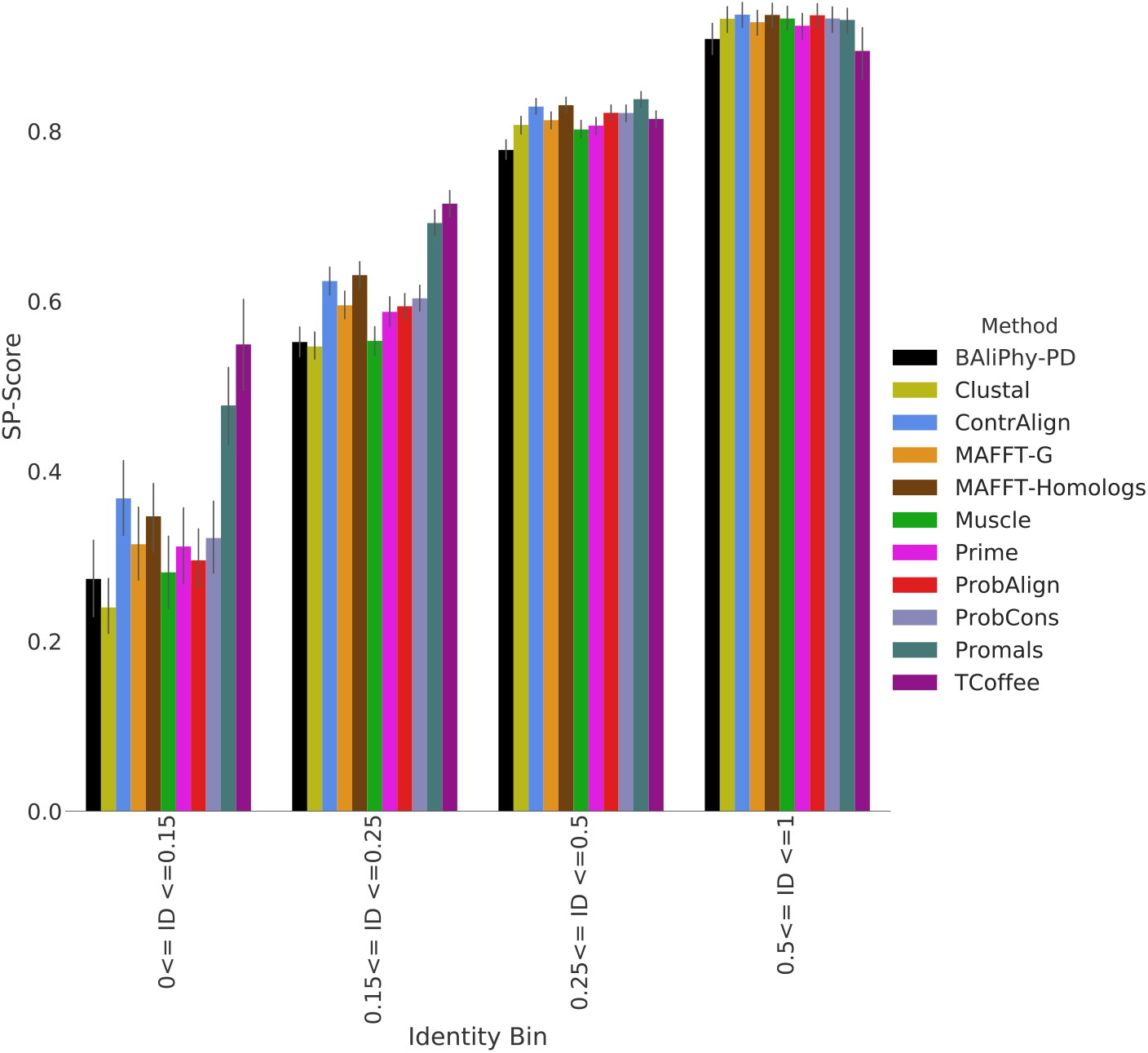
Average SP-Scores for the top methods on the 1192 biological benchmark data sets, binned into different average pairwise sequence identity (ID) levels. The four bins based on average sequence identity, ordered from smallest to largest, have 83, 417, 615, and 77 alignments, respectively.

### Results on Simulated Data Sets

We explored the relative and absolute accuracy of the multiple sequence alignment methods on simulated data sets with 27 sequences. PROMALS and T-Coffee were run on two model conditions (one with high and one with low substitution rates) and had mediocre SP-scores and Modeler scores, clearly neither among the worst nor among the best (see Supplementary Materials). Hence, these methods have poorer accuracy on the simulated data sets than on the biological data sets, a result that is most likely explained by the fact that these methods depend on similarity between the input sequences and those found in external protein benchmark databases, suggesting that simulated amino acid sequences are not very similar to biological amino acid sequences.

The accuracy of the other MSA methods (i.e., MAFFT-G-ins-i, Prank, Prime, Probcons, Probalign, Clustal-Omega, Muscle, and BAli-Phy) varied across these six model conditions, with all methods having the best accuracy for each criterion under the conditions with the lowest substitution and indel rates and the poorest accuracy when both rates were high (Fig. 6; see also Supplementary Materials). When both rates are low, the average PID is low, and all methods had excellent Modeler and SP-scores and the differences between them were small. Thus, the simulation study confirms the trends seen on the biological data sets that average PID impacts accuracy for all MSA methods we explored.

**Figure 6:**
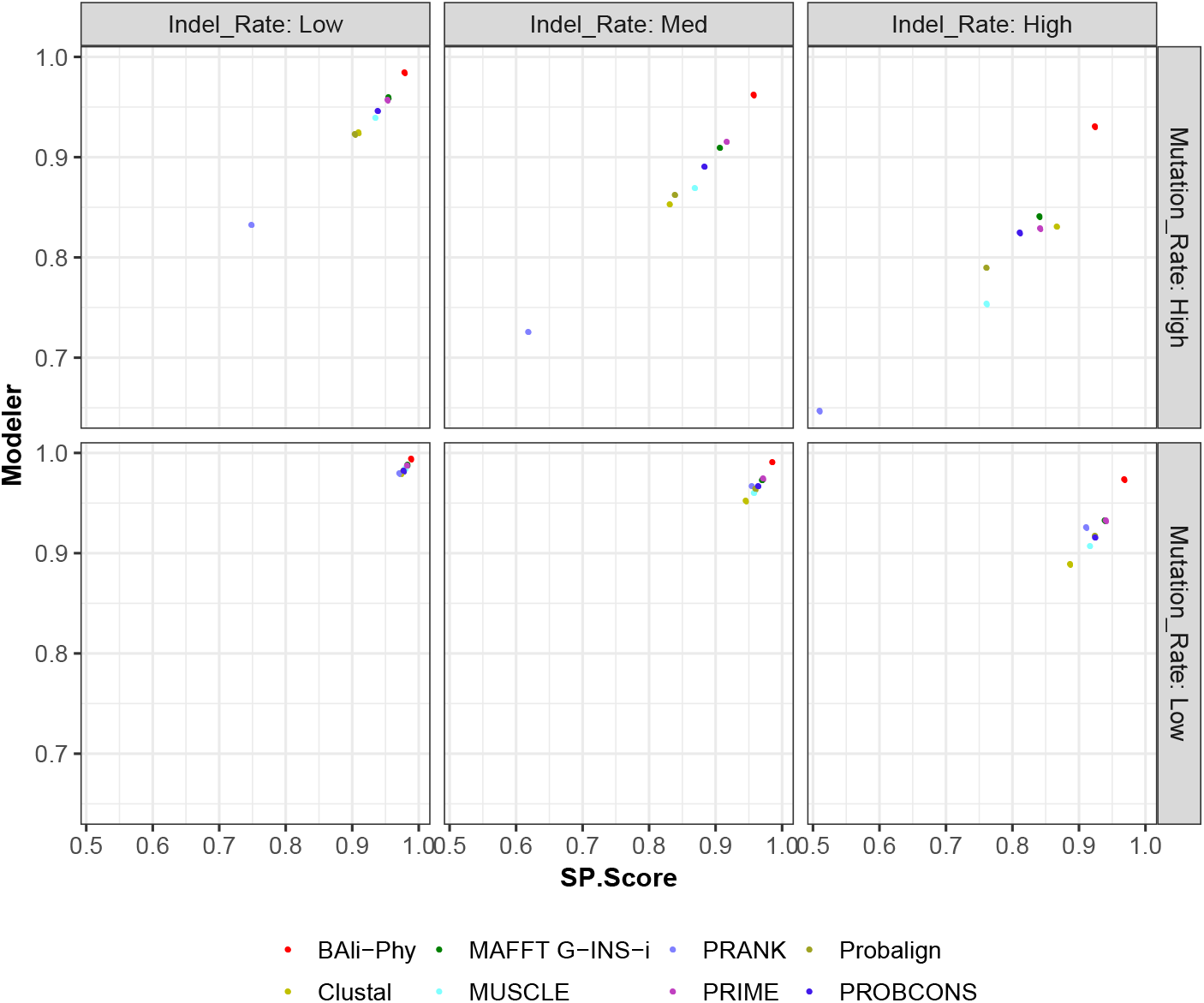
Modeler score vs. SP-Score for MSA methods on simulated amino acid data sets with 27 sequences for 6 different model conditions that vary by the substitution rate and indel rate; averages over 20 replicates are shown.

The most striking observation on the simulated data sets is that BAli-Phy had the best accuracy of all methods with respect to both criteria. Furthermore, while the difference between BAli-Phy and the next best method was small for the easiest model condition, the difference in accuracy increased as the indel rate or the substitution rate increased, and was large under the harder model conditions. For example, under the most difficult model condition (where substitution and indel rates are the highest), BAli-Phy achieved average SP-score and Modeler score of 92-93%, while the second most accurate method had scores that were at least 8% lower (see Supplementary Materials). The relative performance between the other methods depended on the model condition, with Prank having the lowest accuracy when the mutation rate was high, but having reasonable accuracy (falling in the top half of the group) for the low mutation rate conditions, and Clustal having the lowest accuracy for the low mutation rate conditions. Finally, Prime and MAFFT G-INS-i typically came in among the top few methods under all model conditions, but clearly much less accurate than BAli-Phy except for the easiest model conditions where all methods had excellent accuracy.

As shown in Figure 7, similar trends were seen with respect to expansions ratios: BAli-Phy had nearly perfect expansion ratios (i.e., very close to 1.0), whereas most of the other MSA methods (especially Clustal-Omega, MAFFT G-INS-i, and Muscle) often over-aligned, a trend that has been noted before [7, 30]. Probalign had mixed results, sometimes over-aligning and sometimes under-aligning, but not too badly. The outlier here is Prank, which tends to under align (producing expansion ratios greater than 1.0), and in the hardest model condition produced alignments that were 50% longer than the true alignment (see Supplementary Materials).

**Figure 7:**
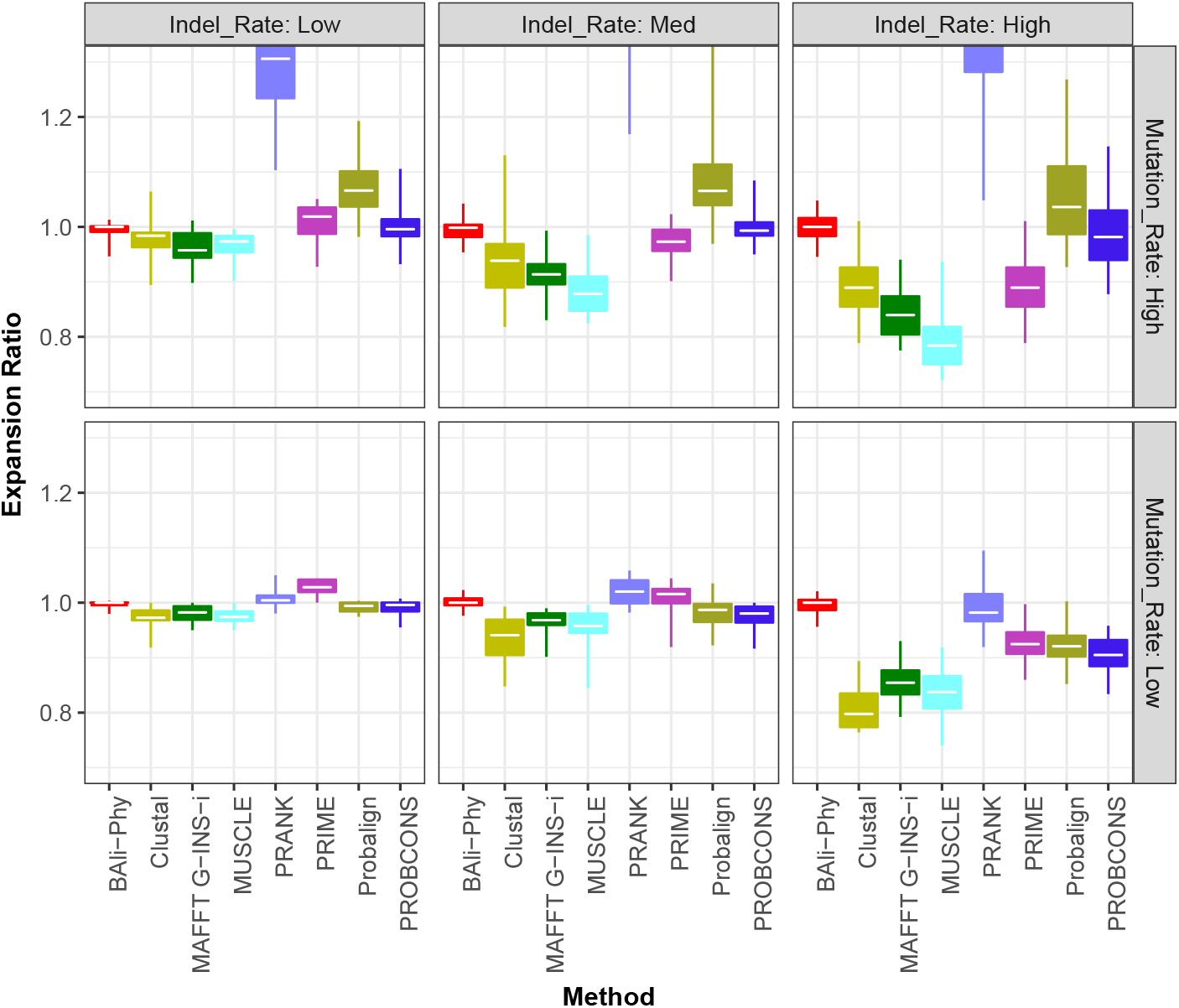
Expansion ratios (1.0 is perfect) for MSA methods on simulated amino acid data sets with 27 sequences for 6 different model conditions that vary by the substitution rate and indel rate; averages over 20 replicates are shown.

#### The impact of model misspecification on BAli-Phy

Finally, we explored the accuracy of BALi-Phy when the protein substitution model it assumes is different from the true substitution model. Our simulation was performed under the WAG substitution model, and so we explored the impact of specifying the true substitution model and a wrong substitution model (JTT) on the resultant alignments by BAli-Phy. As seen in the Supplementary Materials, using the wrong model reduced the alignment accuracy by a very small amount. Under low to moderate rates of evolution, the impact on alignment accuracy is less than 1%, and under the highest rates of evolution that we explored the impact could reach 2%. However, even when affected by model misspecification, BAli-Phy still clearly dominated, by a large margin, the other alignment methods.

### Running Time

The last experiment is to provide an estimate of the running time of different methods. We selected four data sets (one from each of the benchmark collections), each containing 17 sequences to enable this comparison. This comparison is meant to be approximate, as we used different platforms for the methods, and did not ensure that all methods were run using the same environments. T-Coffee and BAli-Phy were run on the National Center of Supercomputing Applications Blue Waters supercomputer and the rest of the methods were run on the Campus Cluster at the University of Illinois at Urbana-Champaign. Some of these methods were compiled from the source code, and we used the precompiled versions for other methods. The running time for BAli-Phy is based on 48 hours for each run, and we ran BAli-Phy 32 independent times.

As shown in Table 3, BAli-Phy is the most computationally intensive of all the methods. T-Coffee and Promals are the next most computationally intensive, followed by Prank. The remaining methods are all reasonably fast, most completing in seconds on the selected data sets.

**Table 3:** Running time information of a single 17-sequence data set in each of the biological benchmarks for different alignment methods, with methods roughly sorted by running time from fastest to slowest. The running times are rounded to the nearest hundredth of a second, and reflect wall clock time. The time reported for most methods is based on a single processor. However, BAli-Phy was run 32 independent times, and the running time reported is for a single run; MAFFT uses 4 threads, and Clustal-Omega uses 12 threads.

| **Benchmark** | **MattBench** | **Homstrad** | **Sisyphus** | **BALiBASE** |
| --- | --- | --- | --- | --- |
| *Data set* | *SF054* | *proteasome* | *AL00048098* | *BALBS213* |
| *Max. Seq. Len.* | *270* | *250* | *117* | *688* |
| DiAlign | 0.0 | 0.0 | 0.0 | 0.0 |
| Prime | 0.1 | 0.0 | 0.0 | 0.0 |
| KAlign | 0.1 | 0.0 | 0.0 | 0.1 |
| Clustal-Omega | 0.4 | 0.3 | 0.1 | 1.5 |
| Muscle | 0.5 | 0.4 | 0.1 | 1.0 |
| MAFFT-G-INS-i | 0.7 | 0.7 | 0.3 | 2.0 |
| ProbAlign | 1.7 | 1.4 | 0.4 | 7.9 |
| ProbCons | 3.1 | 2.6 | 0.6 | 12.6 |
| CONTRAlign | 5.8 | 6.2 | 1.4 | 42.0 |
| Prank | 48.5 | 1:16.1 | 9.4 | 4:14.7 |
| Promals | 14:11.5 | 12:22.1 | 5:06.2 | 24:03.2 |
| T-Coffee | 46:47.2 | 58:04.7 | 7:06.5 | 59:18.8 |
| BALi-Phy | 48:00:00.0 | 48:00:00.0 | 48:00:00.0 | 48:00:00.0 |

## Discussion

The results on the simulated and biological benchmarks are very similar in most respects, but not in all. For both types of data, the best accuracy was obtained for the conditions with the lowest rates of evolution, and the differences between methods were minimal. However, when evolutionary rates were high enough, the differences between methods increased, and some methods clearly outperformed others. Since the MattBench data sets have the lowest average PID, it is not surprising that the alignment methods also demonstrate the lowest average accuracy on MattBench compared to the other benchmarks. Similarly, the Homstrad data sets have the highest average PID of all these benchmarks, and the accuracy was highest on these data sets.

On biological data sets, BAli-Phy had the best Modeler scores and the worst SP-scores across all levels of heterogeneity, while T-Coffee and Promals generally had among the best accuracy (although the relative performance depended on the level of heterogeneity and the criterion). For example, T-Coffee had the best SP-scores for the high heterogeneity data sets (when PID was low) but not under the lowest heterogeneity data sets where Promals and many other methods had better SP-scores. Results on the simulated data sets show different trends: T-Coffee and Promals were not among the better methods on the simulated data sets for either criterion, and BAli-Phy clearly dominated all the other methods with respect to both criteria. Hence, the relative accuracy of methods seems to depend on the heterogeneity in the data set (as measured using PID), the criterion (i.e., Modeler score or SP-score), and – to some extent – whether the data were biological or simulated.

The performance of Prank, a “phylogeny-aware” method that has been referred to as a “heuristic to full statistical alignment” [8], is also worth commenting on. On the biological data sets we examined, Prank has overall among the lowest accuracy of all tested methods. On the simulated data sets, Prank has among the lowest accuracy of the “top performing” methods whenever the substitution rate is high, and is only competitive with the better methods under the lower substitution rates. The poor accuracy on the simulated data sets of Prank under higher rates of evolution is perhaps surprising, given that prior studies have suggested that Prank provides superior alignment accuracy [37]. However, a careful examination of [37] reveals that the simulation conditions in which Prank provided outstanding accuracy had substitutions operating under the simplest model (Jukes-Cantor with a strict molecular clock), which may have favored Prank in some way.

## Conclusions

Statistical sequence alignment, and in particular statistical co-estimation of multiple sequence alignments and phylogenetic trees under stochastic models of sequence evolution that are based on phylogenetic trees, has been considered by many to be the most rigorous approach to alignment estimation.

Our study shows that BAli-Phy, a leading statistical method for co-estimating alignments and trees, has outstanding accuracy on simulated data sets but much lower accuracy on the biological data sets. Specifically, although BAli-Phy often has very good (and sometimes the best) Modeler scores on the biological data, it under-aligns on these datasets, as evidenced by its low SP-scores and high expansions ratios. Put differently, BAli-Phy exhibits both high precision and recall on simulated data but exhibits high precision and low recall on the biological data. Thus, overall accuracy on simulated data and accuracy on biological benchmarks are not necessarily correlated. Most importantly, on the biological data sets, BAli-Phy does not produce alignments with SP-scores that are nearly as good as many popular methods, such as MAFFT and Muscle, that are much faster to use.

This contrast in performance is disturbing, and requires some explanation. There are multiple possible explanations, discussed in detail below, that center on the possible distinctions between evolutionary and structural alignments, and the potential for model misspecification between the model assumed in BAli-Phy and how proteins evolve. Each explanation is likely to be valid, but the relative contribution of each factor is unknown at this time (and beyond the scope of this study). However, some of these factors if they turn out to be significant reasons for this contrast in performance have ramifications in phylogenetics that are important to consider.

One possible explanation is that the reference alignments are accurate evolutionary alignments, but that the sequence evolution model assumed by BAli-Phy is a poor match to the true sequence evolution model under which the proteins evolve. Similar critiques have been made about sequence evolution models used in phylogeny estimation [76, 35] and in simulation studies [28, 9]. Two of the major concerns about these sequence evolution models is the assumption that the sites evolve identically and independently (the *i.i.d.* assumption) and without any selection occurring, which are not realistic for protein sequences. Although the sequence evolution model underlying BAli-Phy is more complex than the standard sequence evolution models discussed in these papers in that it addresses insertions and deletions (i.e., indels) rather than only substitutions, the sequence evolution model nevertheless also has the two problematic features (*iid* site evolution and no selection operating) that were criticized in [76, 35, 28, 9]. Hence, most likely there is substantial model misspecification between the BAli-Phy model of sequence evolution and protein sequence evolution.

If the degree of model misspecification between the model in BAli-Phy and how proteins actually evolve is sufficient to explain much of the distinction in performance between BAli-Phy on biological and simulated datasets, then there are multiple consequences for phylogenetic estimation. Most immediately, if the model misspecification is sufficient to cause protein alignment estimation based on the models to be incorrect, then it suggests the possibility that phylogeny estimation based on these models may be similarly impaired. Hence, better sequence evolution models that more faithfully characterize the evolutionary processes underlying protein sequences will be needed. Furthermore, since many genomic regions (e.g., protein-coding sequences) also evolve under processes that are not *i.i.d.* and that have selective pressures, then model-based phylogeny estimation may also be impaired for many types of markers, at least when based on standard models of sequence evolution. This is the most disturbing of the possible explanations, in terms of the impact on phylogeny estimation.

There are, of course, other potential explanations for the distinction in performance on biological and simulated protein sequences. For example, it is possible that the reference alignments for the biological benchmarks are insufficiently accurate. This might occur is if the reference alignments themselves have false positive homologies (i.e., are over-aligned); in this case, the true alignment would have a high Modeler score and a low SP-score with respect to the reference alignment, which is what we tend to see with BAli-Phy on the biological data sets. If this is the case, then more accurate structural alignments would need to be developed, in order to provide strong and reliable benchmarks. While some error in these reference alignments seem likely, it does not seem very likely that they would be sufficiently incorrect so as to create a condition in which BAli-Phy has much poorer accuracy on biological data than standard alignment methods.

A final possible explanation is that the reference alignments are accurate as structural alignments but not as evolutionary alignments. This is certainly possible, because the distinction between the two types of alignments is real, and the potential for “structural homology” to be different from “evolutionary homology” has been pointed out in several other studies (e.g., [62, 28, 12, 10]). However, it seems unlikely that the differences between correct structural alignments and correct evolutionary alignments would be large enough (and frequent enough) to cause BAli-Phy to consistently be among the least accurate alignment methods in terms of SP-score. Hence, the most likely explanation may be model misspecification between BAli-Phy’s model and how proteins actually evolve, but determining the relative contribution of each of these possible explanations is beyond the scope of this study and is left to future research.

## Data availability

All biological data sets studied in this paper are available in public repositories, and the simulated datasets are available from the authors upon request. The software used to analyze the data sets are also publicly available.

## Supporting information

### Supplementary materials

This document (PDF) has the control file for the simulation study as well as additional discussion.

## Author contributions

Conceived of the project: TW. Designed the experiments: TW MN. Performed the experiments: ES MN. Analyzed the data: TW MN ES. Created figures: MN ES. Wrote the paper: TW MN. All authors read and approved the final manuscript.

Supplementary Materials for: Benchmarking Statistical Alignment Methods

Michael Nute, Ehsan Saleh, and Tandy Warnow The University of Illinois at Urbana-Champaign

## 1 Supplementary Methods

### 1.1 Protocol for Amino Acid Simulations

#### Model Tree Selection

The tree for these simulations was generated by pulling the reference alignment for the protein family Serine Protease from the Homstrad data (filename: `sermam.faa`). The initial goal had been to find a dataset with 25 sequences, but no dataset had exactly that number; the closest dataset (in terms of number of sequences) was `sermam.faa`, which had 27 sequences. We constructed a maximum likelihood tree on the reference alignment for this dataset, using RAxML using the following command:

~~~
<rml>/raxmlHPC-PTHREADS-SSE3 -m PROTGAMMAAUTO -s <aln> -p 12345 -T 12 -n sermam -w./tree
~~~

The tree this yielded is contained in the Indelible control file in the following section.

#### Control file for the simulation

The following block contains the full text of the control file used for these simulations. The entry `<replicate>` on the final line is replaced by the replicate number (0 through 19) prior to running.

~~~
[TYPE] AMINOACID 2
~~~

~~~
[MODEL] MYgtr
[submodel] WAG
[indelmodel] NB 0.637 2
[indelrate] 0.01
[rates] 0 1.0 0
~~~

~~~
[TREE] sermam (((1hcga:0.32813366,1kigh:1.82565029):5.04507020,(1trma:1.15447822,
(1mcta:0.62361724,(2ptn:1.24168792,((2tbs:2.86128857,1a0ja:1.55107647):0.57961993,
(((1ab9:7.47264901,(((1a5ia:1.86915958,1a5ha:0.75672770):5.42205227,
1lmwb:6.73311957):6.89530909,((1sgt:15.58003595,(1bbr:0.57706215,1ppb:0.80065367)
:11.39891397):1.44976925,(1a0la:7.07161115,3est:8.44608968):0.79813103):1.20035927)
:0.00001000):1.10595446,(1dfpa:9.55074028,(((3rp2a:3.98254555,1klt:2.79818431)
:6.79824585,((1hnee:5.51083303,1fuja:2.50912876):1.85070844,1a7s:7.38514536)
:3.64213522):1.00597677,1azza:9.01656013):1.96095053):1.43056606):1.80265480,
((2pka:4.13942226,1ton:4.81015672):3.48654869,1npma:5.54688340):3.87351624)
:1.92885001):0.78867819):0.37891363):0.70651560):6.45102778,1fxya:0.00001000)
:0.0000000);
~~~

~~~
[PARTITIONS] part [sermam MYgtr 200]
~~~

~~~
[SETTINGS]
[output] FASTA
~~~

~~~
[EVOLVE] part 1 R<replicate>
~~~

### 1.2 Sequence sampling protocol

For the biological benchmark datasets, we subsampled 25 sequences from each of the datasets with more than 25 sequences (the “large” alignments). To do this, we selected the number of sequences to sample from 5 up to 25, picking each one in order, and then starting again from 5; thus, 5 sequences were randomly sampled from the first large alignment, 6 sequences from the second large alignment, and so on, until we reached the 22nd dataset where we started with 5 again.

## 2 Additional results

### 2.1 Evidence that BAli-Phy had converged on the biological datasets

The MattBench datasets were the most challenging biological datasets for any method to align, and the ones where BAli-Phy had the worst accuracy. Hence, we report the empirical statistics provided by BAli-Phy that evaluate the evidence that BAli-Phy has converged. BAli-Phy judged 257 of the 259 MattBench datasets to have successfully converged during burn-in, and showed mean minimum ESS values that were greater than 96,000. The Sisyphus datasets were the second hardest; BAli-Phy judged 125 of the 126 Sisyphus dataset to have converged during burn-in, and showed mean minimum ESS values that were greater than 182,000. These statistics suggest that BAli-Phy had successfully converged (at least according to these tests) in analyzing these biological datasets. As noted, the other biological datasets and even the hardest simulated datasets were much easier for BAli-Phy to align, and so there is less need to evaluate convergence on these datasets.

### 2.2 Comparison of T-COFFEE and PROMALS on Simulated Data

Because T-COFFEE and PROMALS rely on retrieval of putative ortholog proteins from public databases as a central component of their alignment algorithm, their alignments of simulated data would not be expected to have high accuracy. Thus, they were not included in the data presented in the main paper. Nonetheless, they were run on the simulated data as a control, and the results are presented here.

Both methods were run on all 20 replicates for both substitution error conditions (trees with scale factor 1.0 and 3.0), each at the original indel rate of 0.01. This has the effect of making the simulations from the 3.0 model tree considerably more difficult.

